# Berberine reverses multidrug resistance in Candida albicans by hijacking the drug efflux pump Mdr1p

**DOI:** 10.1101/2020.06.26.173484

**Authors:** Yaojun Tong, Nuo Sun, Xiangming Wang, Qi Wei, Yu Zhang, Ren Huang, Yingying Pu, Huanqin Dai, Biao Ren, Gang Pei, Fuhang Song, Guoliang Zhu, Xinye Wang, Xuekui Xia, Xiangyin Chen, Lan Jiang, Jingyu Zhang, Liming Ouyang, Buchang Zhang, Yuanying Jiang, Xueting Liu, Richard Calderone, Fan Bai, Lixin Zhang, Gil Alterovitz

## Abstract

Clinical use of antimicrobials faces great challenges from the emergence of multidrug resistant (MDR) pathogens. The overexpression of drug efflux pumps is one of the major contributors to MDR. It is considered as a promising approach to overcome MDR by reversing the function of drug efflux pumps. In the life-threatening fungal pathogen *Candida albicans*, the major facilitator superfamily (MFS) transporter Mdr1p can excrete many structurally unrelated antifungals, leading to multidrug resistance. Here we report a counterintuitive case of reversing multidrug resistance in *C. albicans* by using a natural product berberine to hijack the overexpressed Mdr1p for its own importation. Moreover, we illustrate that the imported berberine accumulates in mitochondria, and compromises the mitochondrial function by impairing mitochondrial membrane potential and mitochondrial Complex I. It results in the selective elimination of Mdr1p overexpressed *C. albicans* cells. Furthermore, we show that berberine treatment can prolong the mean survival time (MST) of mice with a blood-borne dissemination of Mdr1p overexpressed multidrug resistant candidiasis. This study provided a potential direction of novel anti-MDR drug discovery by screening for multidrug efflux pump converters.

## Introduction

Considering the high mortality of fungal infections in immunocompromised patients and the limited number of effective and safe antifungal drugs, development of new antifungals and/or antifungal therapeutics is critical (Gow, van de Veerdonk et al., 2012). Widespread and repeated use of current antifungals, particularly azoles, however, has led to the rapid occurrence of antifungal resistance (Cowen, 2008). Current approaches for novel antifungals discovery are usually targeting essential fungal genes or metabolic pathways, which is very likely to generate drug resistance over time.

One major mechanism underlying fungal drug resistance is the overexpression of drug excretion transporters. There are two types of such transporters in the most commonly seen clinical fungal pathogen *C. albicans*. The *C. albicans* drug-resistance (CDR) transporters, like Cdr1p and Cdr2p, which belong to the ATP-binding cassette (ABC) family, use ATP as their energy source for drug excretion (Holmes, Lin et al., 2008). The other type is the major facilitator superfamily (MFS) transporter, such as Mdr1p (also known as the benomyl/methotrexate resistance protein). This superfamily is a drug / H^+^ antiporter which uses the proton gradient across the cytoplasmic membrane for drug excretion (Pasrija, Banerjee et al., 2007, Yan, 2013, Yan, 2015). Enhanced expression of *MDR1* has been correlated with resistance to a variety of structurally unrelated compounds, such as fluconazole and cerulenin (Hiller, Sanglard et al., 2006). In theory, as a proton gradient driven drug / H^+^ antiporter, Mdr1p has the potential to work reversely, importing substrates while exporting H^+^. This could provide a novel opportunity for antifungals to fight against multidrug resistant *C. albicans*.

Berberine is an alkaloid with a long history of medicinal application in traditional Chinese medicine that can be produced by many plant species, such as *Coptis chinensis* (Coptis, goldenthread), *Hydrastis Canadensis* (goldenseal), and *Berberis vulgaris* (barberry). Berberine has demonstrated significant activities on anti-microbial (Sack & Froehlich, 1982), anti-tumor (Meeran, Katiyar et al., 2008), anti-inflammatory (Kuo, Chi et al., 2004), anti-diabetes (Yin, Xing et al., 2008), lower-cholesterol (Kong, Wei et al., 2004), and compromise-mitochondrial function (Pereira, Branco et al., 2007). Recent studies indicated that fungal mitochondria might be a potential antifungal target due to the presence of unique DNA / proteins (Li & Calderone, 2017). In this study, we described an unexpected association between the killing effect of berberine and the expression of Mdr1p in *C. albicans*. Instead of being excreted, berberine can hijack the overexpressed Mdr1p to facilitate its own accumulation. Berberine then compromises the function of mitochondria to selectively eliminate the Mdr1p overexpressed multidrug resistant *C. albicans*.

## Results

### Berberine susceptibility is inversely correlated with *MDR1* expression in *C. albicans*

In order to search for natural products that can utilize the drug excretion transporter for importation and accumulation, 29 clinical *C. albicans* isolates were used (Table S1) to screen our natural products collection of ∼3,800 pure compounds and ∼ 100,000 crude extracts. Many of these clinical isolates overexpress drug excretion transporters, which results in fluconazole resistance (Fig. 1a). However, we observed a cluster of one set of the fluconazole resistant *C. albicans* isolates with *MDR1* overexpression that showed hyper-sensitivity to berberine (boxed panel of Fig. 1a). Next, we investigated the relationship between Mdr1p overexpression and the hyper-sensitivity of berberine. Another set of *C. albicans* strains with different expression levels of Mdr1p were used, namely, CaS, CaR, CaDEL, and CaCOM (Table S1). CaS, isolated from an AIDS patient G, was proven to be the original fluconazole-susceptible strain with basal Mdr1p expression, while CaR, isolated from the same patient after two years of treatment with fluconazole (Franz, Kelly et al., 1998), was highly resistant to fluconazole, and the dominant contributor of fluconazole resistance was Mdr1p overexpression (Hiller et al., 2006, Wirsching, Michel et al., 2000) (Fig. S1). CaDEL was constructed by *MDR1* deletion of CaR (Wirsching et al., 2000), and CaCOM was constructed by *MDR1* reconstitution into CaDEL (Hiller et al., 2006). *MDR1* expression levels of these four strains were confirmed by quantitative real time PCR (Fig. 1b), and the Mdr1 protein abundance was confirmed by Western blotting (Hiller et al., 2006). From the spot assay results (Fig. 1c), growth of those strains with high levels of Mdr1p (CaR and CaCOM) were inhibited by berberine, while they were uniformly resistant to fluconazole (red boxed panel of Fig. 1c). In contrast, CaS and CaDEL, the two strains with basal expression levels of Mdr1p, showed resistance to berberine but were susceptible to fluconazole (Fig. 1c). These results suggested that berberine has a “<u>s</u>electively <u>e</u>liminate the <u>M</u>dr1p <u>o</u>verexpressed <u>*C*</u>. *albicans*” (SEMOC) property.

**Figure 1.**
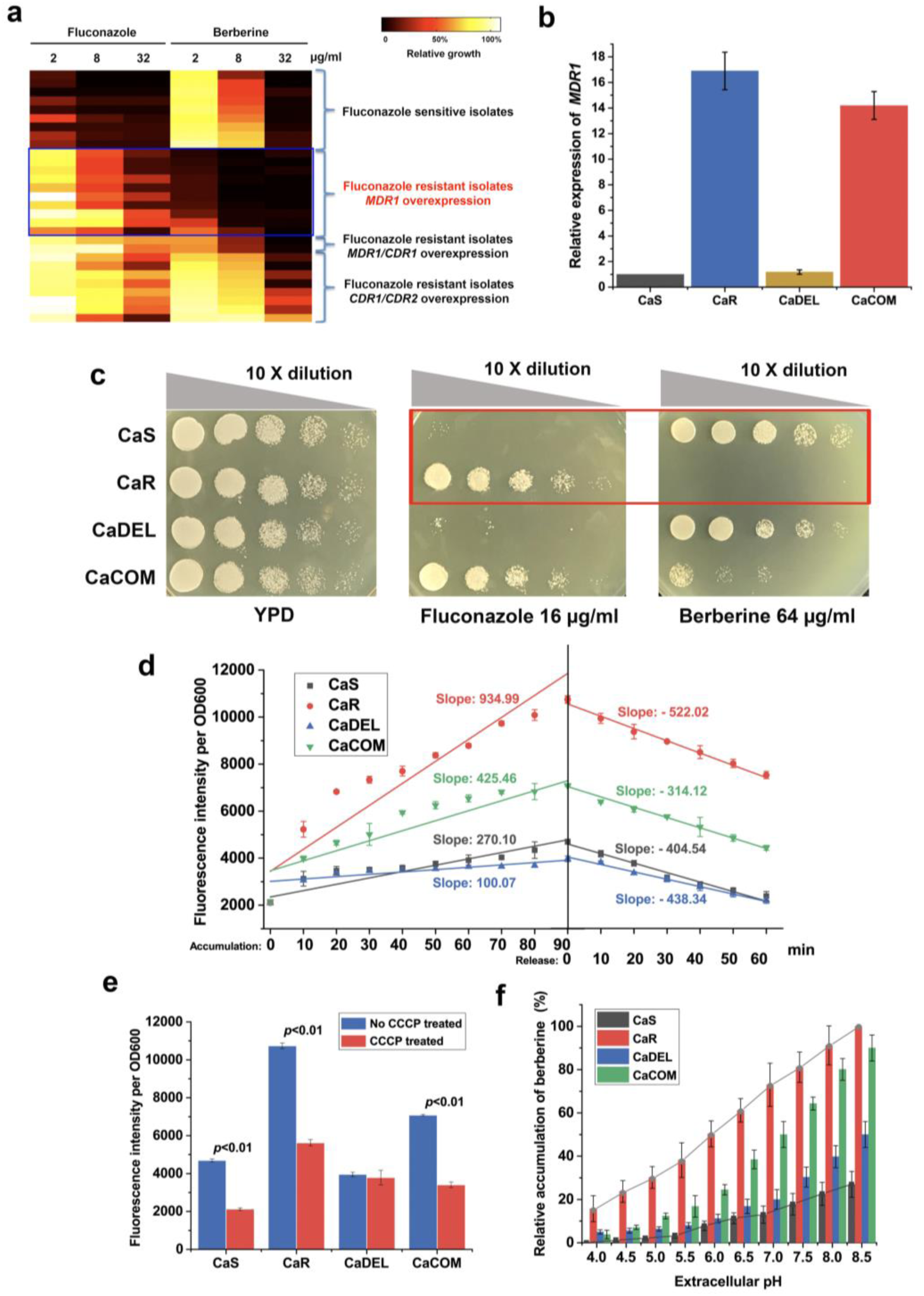
The anti-*C. albicans* activity of berberine is inversely correlated with *MDR1* overexpression. **a**. The antifungal activity of berberine is compared to fluconazole in 29 *C. albicans* isolates, many of which are fluconazole resistant. Susceptibility profiles are indicated as color changes from no growth (black) to growth (white) for each inhibitor (average of three independent experiments). *MDR1* overexpression isolates are hypersusceptible to berberine while resistant to fluconazole. The right panel shows the isolates are clustered according to their susceptibility, source, and/or resistance mechanisms. **b**. *MDR1* expression level of CaS, CaR, CaDEL, and CaCOM, CaS is the original azole-susceptible strain (Franz et al., 1998), while CaR, isolated from the same patient after the long time treatment with fluconazole, is resistant to fluconazole (Hiller et al., 2006, Wirsching et al., 2000), CaDEL is a *MDR1* deletion strain derived from CaR (Wirsching et al., 2000), and CaCOM is the CaDEL strain with reintroduced *MDR1* through expressing the P_*ADH1*_*-MDR1* fusion (Hiller et al., 2006). Quantitative real-time PCR analysis of *MDR1* expression was performed in triplicate. Mean values from three independent experiments are shown. Error bars indicate standard deviation. **c**. Drug susceptibility tested by spot assay. Growth of 10-fold series dilution of CaS, CaR, CaDEL and CaCOM on YPD or YPD containing berberine (64 μg /ml) and fluconazole (16 μg /ml). **d**. A 90-min berberine accumulation time course followed by a 60-min release time course of CaS, CaR, CaDEL and CaCOM. Lines represents the linear fittings, the slope for each line is displayed. **e**. Endpoint accumulation of berberine in CaS, CaR, CaDEL and CaCOM, after a 90-min incubation of berberine, with and without CCCP treatment, respectively. **f**. Extracellular pH effected the berberine accumulation in CaS, CaR, CaDEL and CaCOM.

We speculated that berberine would not be the only natural compound with SEMOC. Besides berberine, other compounds, like jatrorrhizine, proflavine, palmatine and BQM (Sun, Li et al., 2013) which have higher activity against drug resistant *C. albicans* with Mdr1p overexpression over wild type *C. albicans* were also identified from our screening (Fig. S2).

### Berberine could specifically be accumulated in Mdr1p overexpressed *C. albicans* cells

Next, we sought to understand the biological mechanism underlying SEMOC property. As berberine has fluorescence emission at 520 nm by 360 nm excitation, intracellular berberine accumulation can be quantitated by fluorescence readout. We measured the rate of accumulation and release of intracellular berberine in all four strains described in Figure 1b, and 1c, it showed a progressive and consistent increase in berberine accumulation throughout a 90-min incubation period (left panel of Fig. 1d). The amount of berberine in strain CaR is approximately three times as much as that observed in CaS and CaDEL. We observed that the rate of berberine uptake in Mdr1p overexpressed strains exceeded the rate of release (Fig. 1d), leading to a net accumulation of berberine in CaR and CaCOM strains. This observation suggested that overexpressed Mdr1p might serve as an importer of berberine into *C. albicans* cells. To confirm this possibility, we further tested whether the accumulation of berberine was dependent on a proton gradient, as Mdr1p utilizes the proton gradient across the cytoplasmic membrane as its energy source for transportation (Hiller et al., 2006, Pasrija et al., 2007). Carbonyl cyanide m-chlorophenylhydrazone (CCCP) was used to uncouple the proton gradient. Berberine accumulation indeed decreased in Mdr1p-(over)expressed strains after CCCP treatment (Fig. 1e). These data not only confirmed the proton gradient played an important role in berberine accumulation but also indicated that the effects of Mdr1p-overexpression in berberine accumulation.

Due to the drug / H^+^ antiporter property of Mdr1p, we hypothesized that an alkaline extracellular environment might switch Mdr1p from a drug efflux protein into a drug importer. Thus, we evaluated the extracellular pH effect on berberine accumulation. We observed that an increased extracellular pH (reduced extracellular H^+^ concentration) promoted berberine accumulation in strains with a high level of Mdr1p (Fig. 1f).

### Intracellular berberine causes mitochondrial dysfunctions

Then we asked ourselves how intracellular berberine inhibits the growth of *C. albicans*. We reasoned that comparison of the intrinsic differences of response-to-berberine-treatment between CaS and CaR might reveal the targets of berberine. We analyzed the transcriptomes of CaS and CaR with and without berberine treatment, finding that a total of 182 genes were upregulated in CaR (cut-off of 2.0-fold, P-value <0.05 and FDR<0.2) (Fig. 2a), especially genes encoding oxidoreductases (21.7%, GOID: 16491, P-value 5.41×10^−7^) such as the aldo-keto reductase family, *IFD6* and *CSH1*. In order to further narrow down the cellular pathways affected by berberine in *C. albicans*, we performed Gene Set Enrichment Analysis (GSEA), and the ranked gene lists from the transcript profiles were compiled according to the change in their expression to a predefined database of 8,123 gene sets (Uwamahoro, Qu et al., 2012). Significantly enriched gene sets were further visualized using Cytoscape. We observed that the expression of mitochondrial function / aerobic respiration related genes was significantly upregulated after berberine treatment (Fig. 2b and 2c). These results suggested that berberine impacted mitochondrial function.

**Figure 2.**
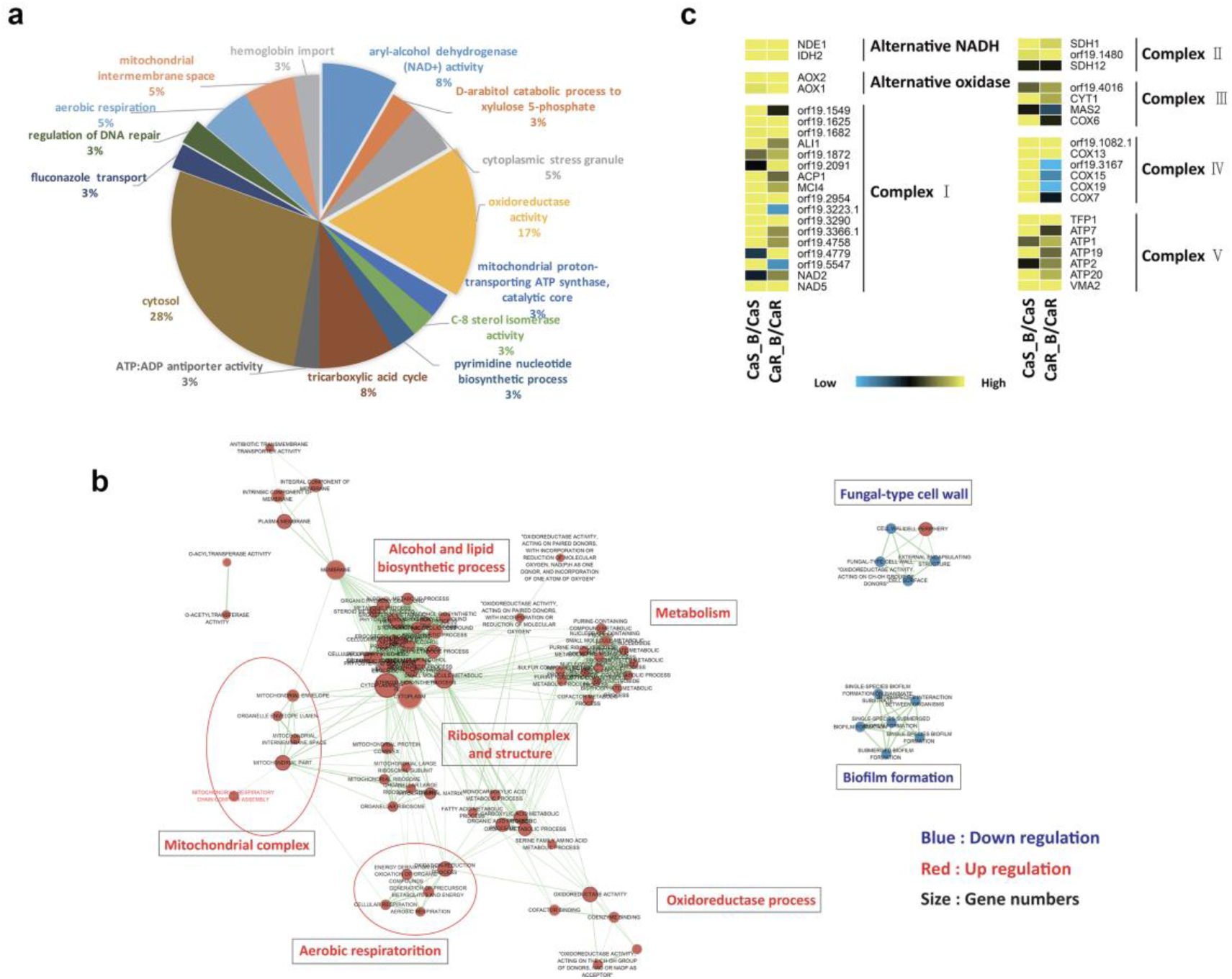
Differential expression analysis of the transcriptome after berberine treatment in CaS and CaR. **a**. RNA-seq analysis of berberine-treated CaR/CaS. Data were presented as a pie chart of functional gene categories (Gene Ontology Term analysis) of upregulated genes in CaR compared to CaS. A total of 182 genes were upregulated in CaR (cut-off of 2.0-fold, P-value <0.05 and FDR<0.2). **b**. The RNA sequencing of the genes differentially expressed after berberine treatment in CaS and CaR; mitochondrial genes were significantly altered. The network of functional groups of genes regulated by berberine was constructed with Cytoscape. Blue circles represent downregulated gene sets, while red circles represent upregulated genes. The size of the circle reflects the number of modulated genes in each functional group. **c**. ETC (Electron Transport Chain) genes were induced by berberine treatment in CaR and CaS. Data were indicated by a blue (downregulation) to yellow (upregulation) growth scale by comparing treated cells with untreated cells of three independent experiments. Genes are clustered by respiratory functional groups. Gene names are taken from the Candida Genome Database (www.candidagenome.org) and their orf19 names are given.

To test this hypothesis, we first confirmed that berberine can accumulate in the mitochondria and as expected, mitochondria of CaR accumulates a higher level of berberine than that of CaS (Fig. 3a). Berberine also exhibited a greater inhibition on non-fermentable carbon sources compared to that on glucose, which possibly indicated that respiration was compromised during berberine treatment (Fig. S3). In addition, important parameters reflecting fungal mitochondrial function were examined. Berberine was found to significantly impair mitochondrial membrane potential (Fig. 3b and Fig. S4), and oxygen consumption (Fig. 3c). Also, after berberine treatment, the activity of the Complex I (NADH dehydrogenase) was sharply reduced (Fig. 3d). In addition, *NDH51*, encoding the mitochondria Complex I 51-kDa subunit of the NADH dehydrogenase protein Ndh51p, was downregulated by 29.9 folds after berberine treatment. To validate Ndh51p as one potential target of berberine, haploinsufficiency (HI) was examined, since the organism is diploid and heterozygote strains lacking one allele usually demonstrate HI. In this regard, the heterozygote *NDH51* mutant demonstrated an HI phenotype that was more susceptible to berberine whereas *ndh51*Δ was more tolerant compared with wild type due to the lack of target gene (Fig. S5). All of these results indicate that berberine interacts with and causes mitochondria dysfunction, which typically stimulates ROS production.

**Figure 3.**
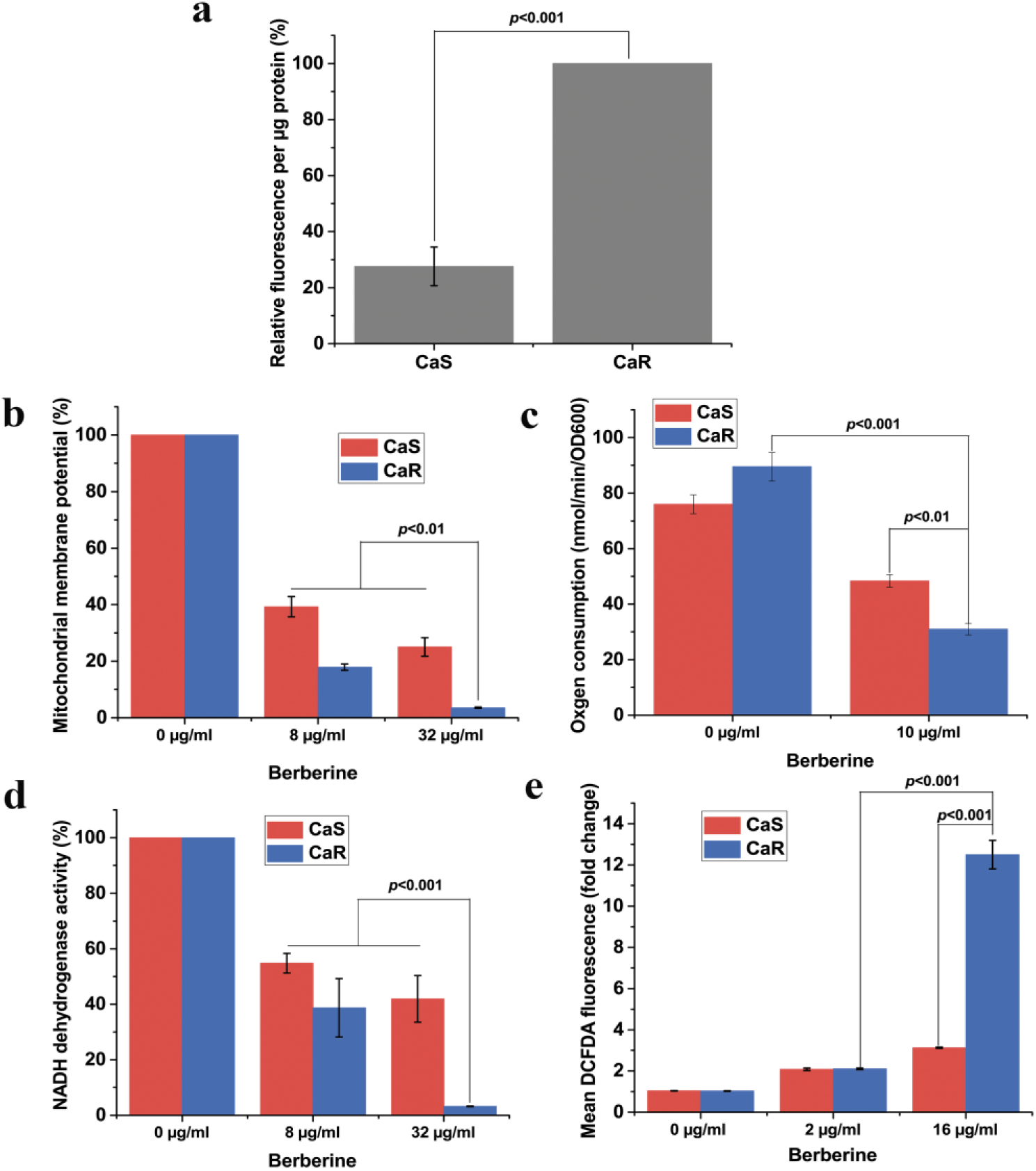
Berberine accumulates in mitochondria and causes mitochondria dysfunction. **a**. Berberine accumulates in mitochondria. Berberine accumulation in percoll gradient purified mitochondria is shown. Fluorescence was measured and normalized to protein concentrations (μg/ml) (mean ± s.d., n=3). The value of CaR was presented as 100%; **b**. Berberine treatment reduces mitochondrial membrane potential (MMP) of CaS and CaR cells. MMP was measured using rhodamine 123 in both strains treated with berberine at indicated concentrations. Fluorescence intensity was monitored with flow cytometry and normalized with control samples. Data are presented as the percentage of control cells (mean ± s.d., n=3); **c**. Berberine inhibits oxygen consumption in *C. albicans*. Respiratory activity of untreated or treated cultures with berberine was measured and normalized to OD_600_ value of cells (mean ± s.d., n=3); **d**. Mitochondrial Complex I (NADH dehydrogenase) activity is inhibited by berberine treatment. NADH dehydrogenase enzymatic activity is normalized to protein content of mitochondria. Data are presented as the percentage of untreated cells (mean ± s.d., n=3); **e**. Berberine induces ROS production. Cells were treated with berberine at the indicated concentrations. ROS levels were measured using 2’,7’-Dichlorodihydrofluorescein diacetate (DCFDA) by flow cytometry and shown as fold-changes (mean ± s.d., n=3).

Consistent with this hypothesis, we found that berberine treatment indeed induced ROS production (Fig. 3e and Fig. S6). In contrast, the addition of antioxidant agents such as ascorbic acid and N-acetyl cysteine (NAC) abrogated inhibitory effects of berberine (Fig. S7). Collectively, these data reveal that berberine activity is related to mitochondrial dysfunction in *C. albicans*. However, we cannot exclude the possibility that berberine, like many other drugs, has multiple targets, given the fact that berberine exhibits a wide spectrum of biological activities.

### Berberine has high potential to be antifungal agent against multidrug resistant invasive fungal pathogens

Besides the novel observation of SEMOC property. We next extensively evaluated the antifungal activity of berberine against several other wild types of common fungal pathogens. We saw that berberine was active against most of those fungal pathogens at relatively low concentrations (from 1 to 16 μg/ml) (Table 1). Remarkably, we also observed that berberine strongly inhibited MDR *Aspergillus fumigatus* (Table 1), the chief cause of invasive aspergillosis (IA). Patients with IA have a mortality rate higher than 90% (Nascimento, Goldman et al., 2003). However, berberine only showed negligible cytotoxicity against several human cell lines (Table 1).

**Table 1.** Berberine is active against various fungal pathogens with low toxicity in human cells.

| Species | Fluconazole | Itraconazole | Berberine |
| --- | --- | --- | --- |
| MIC (µg/ml) |  |  |  |
| *C. albicans* (SC5314) | 0.25 |  | 8 |
| *C. guilliermondii* | 2 |  | 16 |
| *C. glabrata* | 2 |  | 1 |
| *C. tropicalis* | 0.5 |  | 2 |
| *C. parapsilosis* | 1 |  | 16 |
| *C. lusitaniae* | 2 |  | 4 |
| *C. apicola* | 0.25 |  | 4 |
| *C. krusei* | 32 |  | 4 |
| *C. neoformans* (H99) | 4 |  | 4 |
| *C. neoformans* (JEC-21) | 2 |  | 2 |
| *A. fumigatus* (H11-20) |  | 0.5 | 4 |
| *A. fumigatus* (AF293) |  | 0.5 | 4 |
| MDR *A. fumigatus* RIT2 |  | > 100 | 4 |
| MDR *A. fumigatus* RIT3 |  | > 100 | 4 |
| MDR *A. fumigatus* RIT5 |  | > 100 | 4 |
| MDR *A. fumigatus* RIT8 |  | > 100 | 8 |
| MDR *A. fumigatus* RIT10 |  | > 100 | 4 |
| MDR *A. fumigatus* RIT11 |  | > 100 | 4 |
| MDR *A. fumigatus* RIT14 |  | > 100 | 4 |
| HepG2 liver cell |  |  | > 90 |
| NIH/3T3 Fibroblast cell |  |  | > 100 |
| 293T kidney cell |  |  | > 80 |

In order to further evaluate berberine’s potential as an anti-MDR *C. albicans* agent, we tested the antifungal activity of berberine on a set of clinical *C. albicans* isolates, 1#, 4#, 7# and 11#. They were sequentially isolated from an HIV patient who was given an increasing dose of fluconazole during a two years period (White, 1997). As a result, strains 4#, 7# and 11# displayed fluconazole resistance. Interestingly, like CaS and CaR strains, 4#, 7# and 11# strains were also found to have *MDR1* overexpression (by quantitative real time PCR) (Fig. 4a). Consistent with our previous observation, these naturally acquired drug-resistant strains 4#, 7# and 11# showed enhanced susceptibility to berberine over strain 1# (the parental strain) (Fig. 4b). Again, our results demonstrate that the drug resistant *C. albicans* due to Mdr1p overexpression could be specifically inhibited by berberine.

**Figure 4.**
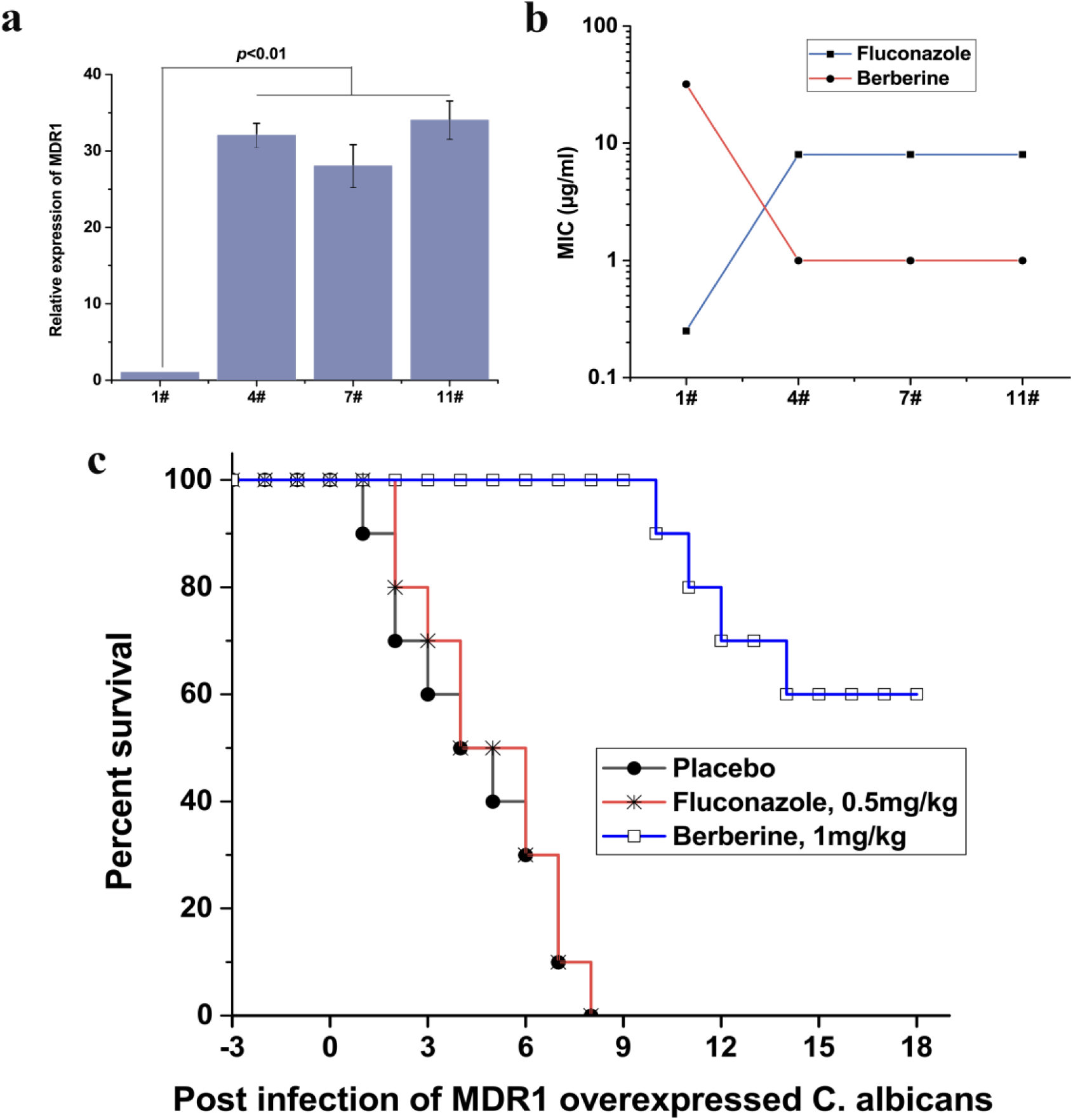
Selective killing of drug-resistant clinical isolates by berberine, both *in vitro* and *in vivo*. ***a***. *MDR1* expression level in *C. albicans* clinical isolates 1# and 4#, 7#, 11# from an HIV patient (White, 1997). Quantitative real time PCR analysis of *MDR1* expression was performed in triplicate. Mean values from three independent experiments are shown. Error bars indicate standard deviation. ***b***. *In vitro* susceptibilities of *C. albicans* isolates 1#, 4#, 7# and 11# to fluconazole and berberine. ***c***. Berberine-treatment dramatically prolonged the life span of the mice that were infected by an Mdr1p overexpressed *C. albicans*.

Candidiasis is often fatal in immunocompromised patients. For this reason, we tested the efficacy of berberine in an immunocompromised animal model of candidiasis. For this purpose, we established an immunocompromised mouse model by intraperitoneal (i.p.) injection of cyclophosphamide (CY) at a dose of 100 mg/kg (body weight) once a day for three consecutive days to specific pathogen-free female ICR mice as described previously (Zhang, Yan et al., 2007). An inoculum (0.1 ml) of 5×10^4^ CFU/ml cells of *C. albicans* strain 11# (fluconazole resistant, while berberine sensitive, *MDR1* overexpression) per mouse killed all mice within 7-8 days (MST, mean survival time, was 4.1 +/- 0.5). Berberine and fluconazole (as a control) were administered by i.p. 6 h post infection and once a day thereafter for three days. A control group of 20 mice received 0.1 ml of diluent (Dulbecco’s phosphate-buffered saline, DPBS) by the same route as the placebo regimens. We saw that berberine treatment dramatically prolonged the MST of those infected mice (p<0.01), while due to the *MDR1* overexpression caused multidrug resistance, fluconazole was failed to save the infected mice (Fig. 4c). From these data, we could conclude that berberine has a very good potential being an antifungal agent against multidrug resistant *C. albicans* due to *MDR1* overexpression.

## Discussion

Multidrug resistance is a worldwide problem that is exacerbated by the shrinking pipeline of new antimicrobial agents. Fungal pathogens adopt intricate strategies to avoid the lethal effects of antibiotics (Anderson, 2005, Cowen, 2008), one of which is overexpression of efflux pump Mdr1p (Hiller et al., 2006). Here we investigated whether Mdr1p, instead of promoting resistance, could be co-opted to promote selective killing of resistant *C. albicans* instead. The enhanced antifungal activity results from increased intracellular accumulation of berberine in *MDR1* overexpressed *C. albicans*. Intriguingly, berberine is reported to accumulate in the rhizome of *Coptis japonica via* an ABC transporter (Cjmdr1p) (Shitan, Bazin et al., 2003), and it can be excreted by some bacterial multidrug excretion transporters, thereby rendering it relatively ineffective as a therapeutic antibacterial agent (Stermitz, Lorenz et al., 2000). Though Mdr1p in *C. albicans*, Cjmdr1p in *Coptis japonica* and NorA (Ball, Casadei et al., 2006, Stermitz et al., 2000) in bacteria do not share structure similarities, they can serve as the channels for berberine, either importing or exporting into cells, which indicated that berberine could be transported by many different transporters. As Mdr1p in *C. albicans* is driven by the proton gradient across the membrane, it has the potential to work bidirectionally. Indeed, we find that increased pH promoted accumulation of berberine, and strains with a high level of Mdr1p expression were more sensitive to pH alteration (Fig. 1f). The findings reported here may represent a novel strategy to overcome MDR, not only in fungal pathogens but also in bacterial pathogens or even other human diseases such as drug resistant cancers. Moreover, our data provide a unique paradigm to explore the function of distinct drug importers. Beyond these observations, a major translational implication of our data is that berberine might make an attractive lead compound for anti-fungal drug development. Finally, we suggest the potential of mitochondria to be a target for new antifungal drug discovery, given that fungal mitochondria have proteins that differ from human mitochondria. Supporting this hypothesis, the deletion of genes encoding these proteins causes cell dysfunction (Li, Chen et al., 2011, Shingu-Vazquez & Traven, 2011, Sun, Fonzi et al., 2013). Our work sheds new mechanistic insights into how membrane transporters facilitate anti-fungal drug resistance. The way berberine hijacks overexpressed transporters to selectively kill drug resistant cells may inspire new strategies for drug discovery and therapeutics to circumvent anti-fungal resistance.

## Materials and Methods

### Strains and growth conditions

The complete list of strains used in this study is listed in Table S1. Fungal strains are stored in 25% glycerol at -80 °C, while cell lines, ordered from cell bank, Shanghai Institutes for Biological Sciences, are stored in liquid nitrogen. RPMI 1640 (Invitrogen) is used according to the manufacturer’s protocol. The YPD medium consisted of yeast extract 1% (w/v), peptone 2% (w/v), and dextrose 2% (w/v), with 2% (w/v) agar to make solid medium when needed.

### Antifungal agents and molecular probes

All chemicals are used according to the manufacturers’ directions. Fluconazole, itraconazole, sanguinarine hydrochloride, jatrorrhizine hydrochloride, palmatine hydrochloride, chelerythrine, and proflavine were purchased from the local chemical pharmacy (purity > 98%). Berberine hydrochloride, cyclophosphamide, rhodamine 123, CCCP (carbonyl cyanide-m-chlorophenyl hydrazone), PI (propidium iodide) and DCFH-DA (2,7-dichlorofluorescein diacetate) were purchased from Sigma-Aldrich (US). Cerulenin was purchased from Alexis, Enzo Life Sciences (US).

### Antifungal susceptibility testing assay

Drug susceptibility testing is carried out as described previously (Zhang et al., 2007) in flat bottom, 96-well microtiter plates (Greiner, Germany), using a broth microdilution protocol modified from the Clinical and Laboratory Standards Institute M-27A methods (CLSI, 2008). MIC is determined as the concentration of drugs that inhibits fungal growth by 80% relative to the corresponding drug-free growth control by reading the optical density (OD600) using a FLUOstar OPTIMA microplate reader (BMG LABTECH, Germany). Spot assay is performed as described previously (Sanglard, Ischer et al., 2003). 10 μl samples of ten-fold serial dilutions of cells, suspended in phosphate-buffered saline (PBS), are spotted onto YPD plates in the absence (control) or in the presence of tested drugs. Photos are taken after a 48-hour incubation at 30 °C. For non-glucose utilization, glucose in yeast extract-peptone agar was replaced with 2% of citrate, glycerol, lactate, or ethanol.

### Berberine accumulation and release assay

All *C. albicans* strains were grown at 30 °C overnight in YPD medium and washed twice with PBS. Cells are resuspended in PBS with approximately 5×10^7^ cells/ml (determined by hemocytometer counting). 32 μg/ml berberine is added into each sample for incubation at 30 °C. For a 90 min accumulation assay, 1 ml of each sample is taken every 10 min and the supernatant is carefully removed by a pipette after 1min of centrifugation at top speed, and then the cell pellets are resuspended in 1ml PBS. For a-60min release assay, 10 ml of each samples at time point of 90 min is centrifuged for 5 min at 5000 g, washed once and then resuspended in 10 ml of PBS. Sampling procedure is similar as accumulation assay. 150 μl samples are transferred into black 96-well microplates with clear bottom (Greiner, Germany) for fluorescence measurements. Fluorescence measurements of berberine are performed with a BioTek™ Synergy™ Mx Monochromator-Based MultiMode Reader (Thermo Fisher Scientific, USA) at 360-nm excitation/520-nm emission wavelengths.

### Accumulation of berberine influenced by CCCP

To investigate the influence of the proton gradient, a final concentration of 20 μg/ml CCCP is added to the samples 30 min prior to the start of accumulation or release assays. Samples were shaken at 200 rpm (30 °C). After incubation, samples are washed with PBS to remove CCCP, then the berberine accumulation assay is carried out as described above.

### pH-dependent accumulation of berberine

To investigate the influence of pH, cells were incubated in RPMI 1640 media adjusted to pH 4.0, 5.0, 5.5, 6.0, 6.5, 7.0, 7.5, 8.0 and 8.5 and treated with berberine. The berberine accumulation assay is carried out as described above.

### RNA preparation, sequencing, and transcriptome analysis

Strains are incubated for 16h at 30 °C in RPMI 1640 medium with shaking with (32 μg/ml) or without berberine. Two volumes of RNAprotect Reagent (Qiagen, Germany) are added into the cultures according to manufacturer’s protocol. Cells are harvested, washed twice and then resuspended into PBS. The cells are then lysed by lysozyme (400 μg/ml, Ready-Lyse™ Lysozyme Solution, Epicenter). Total RNA is purified by RNeasy Mini Kit (Qiagen, Germany). Ribosomal RNA is removed by RiboMinus Transcriptome Isolation Kit (Invitrogen, USA) and mRNA is cleaned up using RNeasy Mini Spin columns (Qiagen, Germany). The resulting mRNA is used for library construction for RNA-seq using NEBNext mRNA Library Prep Reagent set for Illumina (NEB, USA). Briefly, mRNA is fragmented to desired length and reverse transcribed into first strand cDNA. The single strand cDNA is used for synthesizing double strand DNA followed by end repair, dA-tailing, adaptor ligation and PCR amplification. The library is tested by length determination and quantitative PCR quantification. These constructed libraries are then sequenced by Illumina platform by paired-end chemistry.

RNA reads are aligned to the *C. albicans* SC5314 assembly using Tophat (Trapnell, Pachter et al., 2009), and quantified with HTseq (Anders, Pyl et al., 2015). The raw read counts are normalized with DESeq2 (Love, Huber et al., 2014) to estimate gene expression and identify differential gene expression. Differential gene expression is identified using the threshold of the parametric p-value <0.05 and a fold change of at least 2 and FDR < 0.2. Gene ontology analysis is performed at the *Candida* genome database (CGD, www.candidagenome.org) and FungiFun2 (https://elbe.hki-jena.de/fungifun/). Enrichment maps are constructed with Cytoscape 2.8.3 (http://www.cytoscape.org) and the Enrichment Map v1.2 plug-in using the default settings (http://www.baderlab.org/Software).

### Assay of the ROS measurement

As previously described (Li et al., 2011), intracellular ROS production is detected by staining cells with the ROS-sensitive fluorescent dye DCFH-DA (2,7-dichlorofluorescein diacetate; Sigma). Cells from 25 ml cultures grown at 30 °C overnight in YPD medium are collected and washed twice with PBS. The pellets are suspended to 10^6^ cells in 10 ml of PBS plus 2% glucose and treated with or without 10 μM DCFH-DA for 30 min at 37 °C in dark. Cells from each sample are collected and washed twice with PBS after staining. The cells are resuspended with PBS plus 2% glucose and treated with berberine at the indicated concentrations for 60 min 30 °C. The fluorescent intensity is measured using a FACScan flow cytometer (Becton Dickinson). Propidium iodide (PI) was added to each sample to detect dead cells. The mean fluorescence for ROS was quantified only in live cells.

### Assay of protoplast and mitochondrial preparations

Cells are grown in 250 ml of YPD broth overnight at 30 °C, then washed and resuspended in RPMI 1640 medium. After incubation with or without 20 µg/ml berberine for 1 h, cells are harvested by centrifugation (5000 rpm for 10 min) and washed with 50 ml of cold water and buffer A (1 M sorbitol, 10 mM MgCl_2_, 50 mM Tris-HCl [pH 7.8]). Protoplasts are made by digesting the cell wall with Zymolase 20T for 1h at 30 °C. The mitochondrial fraction is obtained by Percoll-density-gradient centrifugation as described previously (Li et al., 2011), and suspended in 1 ml of buffer D (0.6 M mannitol, 10 mM Tris-HCl, [pH 7.0]). Protein content is determined by the Biuret method.

### Assay for the measurement of mitochondrial membrane potential

Mitochondrial membrane potential is determined by staining cells with rhodamine 123 (Ferlini & Scambia, 2007). Cells from 25 ml cultures grown at 30 °C overnight in YPD medium are collected and washed twice with RPMI 1640 medium. The pellets are resuspended in RPMI 1640 medium to 10^6^ cells/ml. Berberine is added at the indicated concentrations. After 60 min incubation, the samples are treated with or without 10 μg/ml rhodamine 123 for 15 min at 37 °C in dark. Cell fluorescence in the absence of rhodamine 123 is used to verify that background fluorescence is similar among strains. Cells from each sample are collected and washed twice with PBS after staining. Fluorescence is measured using a FACScan flow cytometer (Becton Dickinson).

### Assay for oxygen consumption rate

Oxygen consumption is measured polarographically using a Clark-type electrode (Hansatech Instruments). Cells are grown overnight at 30 °C in 20 ml of YPD broth and diluted in fresh YPD broth next day for an additional 4 h until exponential growth is achieved. Cells are then centrifuged, washed with PBS, resuspended in RPMI 1640, and then treated with 10 µg/ml berberine for 2 h. Cells are collected, washed with PBS, and resuspended in 2 ml of YPD broth before loading into a sealed 1.5-ml glass chamber. The oxygen concentration in the chamber is monitored over 5-10 min period. The respiratory rate is calculated as the consumption of oxygen per min per ml of cell suspension normalized by OD_600_ value.

### Complex I (NADH: ubiquinone oxidoreductase) activity assay

Crude mitochondrial preparations are first treated with two cycles of freeze-thawing in a hypotonic solution (25 mM K_2_ HPO_4_ [pH 7.2], 5 mM MgCl_2_), followed by a hypotonic shock in H_2_O. A total of 20 μg of mitochondrial protein from each sample is used to measure complex I enzymatic activity. Mitochondria in 0.8 ml of H_2_O were incubated for 2 min at 37 °C and then mixed with 0.2 ml of a solution containing 50 mM Tris, pH 8.0, 5 mg/ml BSA, 0.24 mM KCN, 4 μM antimycin A, and 0.8 mM NADH (substrate of Complex I). The reaction is initiated by introducing an electron acceptor, 50 μM DB (2,3-dimethoxy-5-methyl-6-n-decyl-1,4-benzoquinone). Enzyme activity is followed as a decrease in absorbance of NADH at 340 nm minus that at 380 nm.

### Relative quantification of differentially expressed genes by quantitative real time PCR

All primers used in this study for qRT-PCR are listed in Table S2. RNA isolation, cDNA synthesis, and PCR amplification are carried out as described previously (Xu, Zhang et al., 2006). Triplicate independent real time PCRs are performed using the Light Cycler System (Roche diagnostics). The change in fluorescence of SYBR Green I in every cycle was monitored by the system software, and the threshold cycle (CT) is measured. 18S rDNA is used as an internal control, and the relative gene expression level is calculated using the formula 2^−ΔΔCT^.

### Murine model of systemic infection and drug treatment

Specific-pathogen-free female ICR [Crl: CD-1] mice (female, white, about 20-22 g) are used throughout the experiment, and the experiments are carried out as described previously (Zhang et al., 2007). Before infection, mice are rendered neutropenic by i.p. injection of cyclophosphamide (CY) (Sigma) daily for three consecutive days at a dosage of 100 mg/kg body weight before infection. Mice are monitored at designated days after the first CY injection for WBC counts by using a hemocytometer. Mice are then infected with 0.1 ml of 5 ×10^4^ CFU/ml cells of *C. albicans* 11# per mouse in warmed saline (35 °C) by the lateral tail vein on day 3 after pretreatment with CY. Berberine (1 mg/kg) and fluconazole (0.5 mg/kg) independently are administered intraperitoneally (i.p.) 6 hours post infection and once daily thereafter for three days. And 0.1 ml of diluent (Dulbecco’s phosphate-buffered saline, DPBS) by the same route as the placebo regimens. Data is averaged from three experiments.

### Cytotoxicity evaluation using MTT assay

The protocol is modified from (Bahuguna, Khan et al., 2017). Briefly, the testing cells were seeded in a 96-well flat-bottom microtiter plate with 1 × 10^4^ cells/well and allowed to adhere for 24 hours at 37 °C in a CO_2_ incubator. Cells are gently washed with fresh medium. Cells are then treated with various concentrations of the target compounds for 24 hours in the same cultivation condition. Cells are gently washed with fresh medium again. Subsequently, 10 μl of MTT working solution (5 mg/ml in phosphate buffer solution) is added to each well and the plate is incubated for 4 hours at 37 °C in a CO_2_ incubator. The medium is then aspirated, and the formed formazan crystals are solubilized by adding 50 μl of DMSO per well for 30 min at 37 °C in a CO_2_ incubator. Finally, the intensity of the dissolved formazan crystals (purple color) is quantified using the ELISA plate reader at 540 nm.

## Data Availability

All data supporting the findings of this study are available within this article and Supporting Information or from the corresponding author on reasonable request.

## Acknowledgments

We thank Joachim Morschhäuser, Dominique Sanglard and Theodore C. White for kindly providing drug-resistant clinical fungal isolates. We thank Haian Fu, Bruce Alberts, Rong An, Elizabeth Ashforth and Glen Bulmer for their critical reading of the manuscript and helpful discussions. This work was supported by the National Natural Science Foundation of China #31720103901, the “111” Project of China #B18022, the Fundamental Research Funds for the Central Universities #22221818014S, the Open Project Funding of the State Key Laboratory of Bioreactor Engineering, and the Shandong Taishan Scholar Award to L.Z. Y.T. was also partially supported by the Novo Nordisk Foundation (NNF10CC1016517).

## Author Contributions

Y.T., N.S., F.B., and L.Z. conceived and designed the experiments. Y.T., N.S., X. W., Y. P., Q.W., B.R., F.S., G.Z., X.W., X.X., X.C., L.J., J.Z., L.O., Y.Z., G.P., Y.P., H.D., and W.F., performed the experiments. N.S., Y.T., B.Z., R.H., X.L., Y.J., G.A., and F.B. analyzed the data. Y.T., F.B., and L.Z. wrote the paper.

## References

Anders S, Pyl PT, Huber W (2015) HTSeq--a Python framework to work with high-throughput sequencing data. Bioinformatics 31: 166–9

Anderson JB (2005) Evolution of antifungal-drug resistance: Mechanisms and pathogen fitness. Nature Reviews Microbiology 3: 547–556

Bahuguna A, Khan I, Bajpai VK, Kang SC (2017) MTT assay to evaluate the cytotoxic potential of a drug. Bangladesh Journal of Pharmacology 12: 115–118

Ball AR, Casadei G, Samosorn S, Bremner JB, Ausubel FM, Moy TI, Lewis K (2006) Conjugating berberine to a multidrug efflux pump inhibitor creates an effective antimicrobial. ACS Chem Biol 1: 594–600

CLSI (2008) In Reference method for broth dilution antifungal susceptibility testing of yeasts; Approved standard, 3rd ed, CLSI document M27-A3, Wayne, PA: National Committee for Clinical Laboratory Standards

Cowen LE (2008) The evolution of fungal drug resistance: modulating the trajectory from genotype to phenotype. Nature Reviews Microbiology 6: 187–198

Ferlini C, Scambia G (2007) Assay for apoptosis using the mitochondrial probes, Rhodamine123 and 10-N-nonyl acridine orange. Nat Protoc 2: 3111–4

Franz R, Kelly SL, Lamb DC, Kelly DE, Ruhnke M, Morschhauser J (1998) Multiple molecular mechanisms contribute to a stepwise development of fluconazole resistance in clinical Candida albicans strains. Antimicrob Agents Chemother 42: 3065–72

Gow NA, van de Veerdonk FL, Brown AJ, Netea MG (2012) Candida albicans morphogenesis and host defence: discriminating invasion from colonization. Nat Rev Microbiol 10: 112–22

Hiller D, Sanglard D, Morschhauser J (2006) Overexpression of the MDR1 gene is sufficient to confer increased resistance to toxic compounds in Candida albicans. Antimicrob Agents Chemother 50: 1365–71

Holmes AR, Lin YH, Niimi K, Lamping E, Keniya M, Niimi M, Tanabe K, Monk BC, Cannon RD (2008) ABC Transporter Cdr1p Contributes More than Cdr2p Does to Fluconazole Efflux in Fluconazole-Resistant Candida albicans Clinical Isolates. Antimicrobial Agents and Chemotherapy 52: 3851–3862

Kong W, Wei J, Abidi P, Lin M, Inaba S, Li C, Wang Y, Wang Z, Si S, Pan H, Wang S, Wu J, Li Z, Liu J, Jiang JD (2004) Berberine is a novel cholesterol-lowering drug working through a unique mechanism distinct from statins. Nat Med 10: 1344–51

Kuo CL, Chi CW, Liu TY (2004) The anti-inflammatory potential of berberine in vitro and in vivo. Cancer Lett 203: 127–37

Li D, Chen H, Florentino A, Alex D, Sikorski P, Fonzi WA, Calderone R (2011) Enzymatic dysfunction of mitochondrial complex I of the Candida albicans goa1 mutant is associated with increased reactive oxidants and cell death. Eukaryot Cell 10: 672–82

Li DM, Calderone R (2017) Exploiting mitochondria as targets for the development of new antifungals. Virulence 8: 159–168

Love MI, Huber W, Anders S (2014) Moderated estimation of fold change and dispersion for RNA-seq data with DESeq2. Genome Biol 15: 550

Meeran SM, Katiyar S, Katiyar SK (2008) Berberine-induced apoptosis in human prostate cancer cells is initiated by reactive oxygen species generation. Toxicol Appl Pharmacol

Nascimento AM, Goldman GH, Park S, Marras SA, Delmas G, Oza U, Lolans K, Dudley MN, Mann PA, Perlin DS (2003) Multiple resistance mechanisms among Aspergillus fumigatus mutants with high-level resistance to itraconazole. Antimicrob Agents Chemother 47: 1719–26

Pasrija R, Banerjee D, Prasad R (2007) Structure and function analysis of CaMdr1p, a major facilitator superfamily antifungal efflux transporter protein of Candida albicans: identification of amino acid residues critical for drug/H+ transport. Eukaryot Cell 6: 443–53

Pereira GC, Branco AF, Matos JA, Pereira SL, Parke D, Perkins EL, Serafim TL, Sardao VA, Santos MS, Moreno AJ, Holy J, Oliveira PJ (2007) Mitochondrially targeted effects of berberine [Natural Yellow 18, 5,6-dihydro-9,10-dimethoxybenzo(g)-1,3-benzodioxolo(5,6-a) quinolizinium] on K1735-M2 mouse melanoma cells: comparison with direct effects on isolated mitochondrial fractions. J Pharmacol Exp Ther 323: 636–49

Sack RB, Froehlich JL (1982) Berberine inhibits intestinal secretory response of Vibrio cholerae and Escherichia coli enterotoxins. Infect Immun 35: 471–5

Sanglard D, Ischer F, Parkinson T, Falconer D, Bille J (2003) Candida albicans mutations in the ergosterol biosynthetic pathway and resistance to several antifungal agents. Antimicrobial agents and chemotherapy 47: 2404–12

Shingu-Vazquez M, Traven A (2011) Mitochondria and fungal pathogenesis: drug tolerance, virulence, and potential for antifungal therapy. Eukaryot Cell 10: 1376–83

Shitan N, Bazin I, Dan K, Obata K, Kigawa K, Ueda K, Sato F, Forestier C, Yazaki K (2003) Involvement of CjMDR1, a plant multidrug-resistance-type ATP-binding cassette protein, in alkaloid transport in Coptis japonica. Proc Natl Acad Sci U S A 100: 751–6

Stermitz FR, Lorenz P, Tawara JN, Zenewicz LA, Lewis K (2000) Synergy in a medicinal plant: antimicrobial action of berberine potentiated by 5’-methoxyhydnocarpin, a multidrug pump inhibitor. Proc Natl Acad Sci U S A 97: 1433–7

Sun N, Fonzi W, Chen H, She X, Zhang L, Calderone R (2013) Azole Susceptibility and Transcriptome Profiling in Candida albicans Mitochondrial Electron Transport Chain Complex I Mutants. Antimicrob Agents Chemother 57: 532–42

Sun N, Li D, Fonzi W, Li X, Zhang L, Calderone R (2013) Multidrug-resistant transporter mdr1p-mediated uptake of a novel antifungal compound. Antimicrob Agents Chemother 57: 5931–9

Trapnell C, Pachter L, Salzberg SL (2009) TopHat: discovering splice junctions with RNA-Seq. Bioinformatics 25: 1105–11

Uwamahoro N, Qu Y, Jelicic B, Lo TL, Beaurepaire C, Bantun F, Quenault T, Boag PR, Ramm G, Callaghan J, Beilharz TH, Nantel A, Peleg AY, Traven A (2012) The functions of Mediator in Candida albicans support a role in shaping species-specific gene expression. PLoS Genet 8: e1002613

White TC (1997) Increased mRNA levels of ERG16, CDR, and MDR1 correlate with increases in azole resistance in Candida albicans isolates from a patient infected with human immunodeficiency virus. Antimicrob Agents Chemother 41: 1482–7

Wirsching S, Michel S, Morschhauser J (2000) Targeted gene disruption in Candida albicans wild-type strains: the role of the MDR1 gene in fluconazole resistance of clinical Candida albicans isolates. Mol Microbiol 36: 856–65

Xu Z, Zhang LX, Zhang JD, Cao YB, Yu YY, Wang DJ, Ying K, Chen WS, Jiang YY (2006) cDNA microarray analysis of differential gene expression and regulation in clinically drug-resistant isolates of Candida albicans from bone marrow transplanted patients. Int J Med Microbiol 296: 421–34

Yan N (2013) Structural advances for the major facilitator superfamily (MFS) transporters. Trends Biochem Sci 38: 151–9

Yan N (2015) Structural Biology of the Major Facilitator Superfamily Transporters. Annu Rev Biophys 44: 257–83

Yin J, Xing H, Ye J (2008) Efficacy of berberine in patients with type 2 diabetes mellitus. Metabolism 57: 712–7

Zhang L, Yan K, Zhang Y, Huang R, Bian J, Zheng C, Sun H, Chen Z, Sun N, An R, Min F, Zhao W, Zhuo Y, You J, Song Y, Yu Z, Liu Z, Yang K, Gao H, Dai H et al. (2007) High-throughput synergy screening identifies microbial metabolites as combination agents for the treatment of fungal infections. Proceedings of the National Academy of Sciences of the United States of America 104: 4606–11

